# Convergent Cerebrospinal Fluid Proteomes and Metabolic Ontologies in Humans and Animal Models of Rett Syndrome

**DOI:** 10.1101/2021.11.30.470580

**Authors:** Stephanie A. Zlatic, Duc Duong, Kamal K.E. Gadalla, Brenda Murage, Lingyan Ping, Ruth Shah, Omar Khwaja, Lindsay C. Swanson, Mustafa Sahin, Sruti Rayaprolu, Prateek Kumar, Srikant Rangaraju, Adrian Bird, Daniel Tarquinio, Stuart Cobb, Victor Faundez

## Abstract

MECP2 loss-of-function mutations cause Rett syndrome, a disorder that results from a disrupted brain transcriptome. How these transcriptional defects are decoded into a disease proteome remains unknown. We studied the proteome in Rett syndrome cerebrospinal fluid (CSF) across vertebrates. We identified a consensus proteome and ontological categories shared across Rett syndrome cerebrospinal fluid (CSF) from three species, including humans. Rett CSF proteomes enriched proteins annotated to HDL lipoproteins, complement, mitochondria, citrate/pyruvate metabolism, as well as synapse compartments. We used these prioritized and shared ontologies to select analytes for orthogonal quantification. These studies independently validated our proteome and ontologies. Ontologically selected CSF hits had genotypic discriminatory capacity as determined by Receiver Operating Characteristic (ROC) analysis and distinguished Rett from a related neurodevelopmental disorder, CDKL5 deficiency disorder. We propose that *Mecp2* mutant CSF proteomes and ontologies inform novel putative mechanisms and biomarkers of disease. We suggest that Rett syndrome is a metabolic disorder impacting synapse function.

## Introduction

The cellular and molecular understanding of neurodevelopmental disorders has been greatly advanced by the study of single gene defects ^1,2^. Among these monogenic neurodevelopmental disorders, Rett syndrome, caused by mutations in *MECP2*, stands out because of its severity and developmental regression. The molecular function of MeCP2 as an epigenetic chromatin regulator is well defined ^3^, affecting the expression of a vast number of RNAs in the brain ^3-6^. The molecular complexity of Rett syndrome is compounded by the extensive and varied modifications of coding and non-coding transcriptomes across brain cell types and regions ^7-11^. This fact makes transcriptional prediction of Rett syndrome proteomes a complex and uncertain endeavor. Thus, we focus on the proteome to identify biochemical and cellular alterations in Rett syndrome brains ^12^. Moreover, known Rett syndrome phenotypes such as synaptic, circuit, and behavioral alterations ultimately originate in alterations of protein expression and function ^4,13,14^. Thus, the proteome decodes *Mecp2*-dependent transcriptional changes and executes a diseased phenome. Despite these advantages of the proteome to illuminate pathogenic mechanisms and to identify disease biomarkers, few studies examine how the proteome is modified in Rett syndrome brains ^15,16^.

Our goal was to identify a Rett proteome capable of distinguishing normal and disease brain states across neurodevelopment that is clinically accessible for sample collection, diagnostic testing, and assessing treatment outcomes. We focused on CSF as a clinically accessible sample whose composition is dictated by the brain and neurodevelopment ^17-19^. CSF carries neurodevelopmental instructive signals and nutrients and accrues secretions and metabolites that reflect functional states of diverse cell types in brain parenchyma and the choroid ^20^. For example, composition of the CSF proteome in individuals with Alzheimer’s predicts key diagnostic molecular pathology in the Alzheimer’s brain ^21-23^. Thus, the CSF proteome has the potential to inform us about brain-wide normal and pathological states.

The study of the CSF in genetic or sporadic forms of neurodevelopmental disorders lags behind similar studies in neurological diseases. Less than a handful of studies analyze the proteome of this biofluid ^24^. The study of the CSF in neurodevelopmental disorders has mostly been focused to targeted studies of few analytes such as cytokines, growth factors, neuropeptides, or metabolites ^25-30^. Thus, we do not know if a neurodevelopmental disorder, such as Rett syndrome, reproducibly and distinctively modifies the CSF proteome to predict disease mechanisms and biomarkers of disease. Here, we address this fundamental question by comprehensively and unbiasedly exploring the CSF proteome of three species carrying mutations in *MECP2*/*Mecp2*. We defined a consensus proteome and ontologies predictive of Rett syndrome disease mechanisms. Analytes found in the *Mecp2*-sensitive proteome behaved with sufficient sensitivity and specificity to act as Rett syndrome biomarkers and to discriminate Rett syndrome CSF from a CDKL5 deficiency disorder CSF, a phenotypically related syndrome ^31-33^. This is the first multispecies study of the CSF proteome in a monogenic disorder of neurodevelopment. Based on our proteomic analysis, we propose that Rett syndrome is a synaptic and metabolic neurodevelopmental disorder. Further, our experimental strategy offers a platform for the identification of proteomes and biomarkers in the CSF of any childhood genetic neurological disorder and to infer putative mechanisms of disease.

## Results

The CSF proteome is composed of proteins secreted by conventional and non-conventional secretory pathways, exosomes, and ectosomes from brain ^34^. We collectively refer to these proteins as the brain secreted proteome. We sought to identify secreted proteins sensitive to MECP2 gene defects. To achieve this goal, we designed a multipronged strategy to quantify secreted proteomes from wild type and *MECP2*-null neuron conditioned media, cerebrospinal fluid from *Mecp2* null male rat and mouse models, and the CSF from female individuals with Rett syndrome collected before and after recombinant IGF1 treatment (Fig. 1A) ^35,36^. We reasoned that overlapping proteins and ontologies across diverse experimental systems would identify robust proteins and ontologies to inform putative disease mechanisms and Rett syndrome biomarkers. Furthermore, we designed our studies with an emphasis in replicability across experimental sites, quantification platforms, and species to inform biomarker selection (Fig. S1). We chose to quantify proteomes with Tandem Mass Tagging (TMT) mass spectrometry as a high precision method ^37,38^. TMT datasets were analyzed by fold of change/p value volcano thresholding plus machine learning approaches (Fig. 1A).

**Figure 1.**
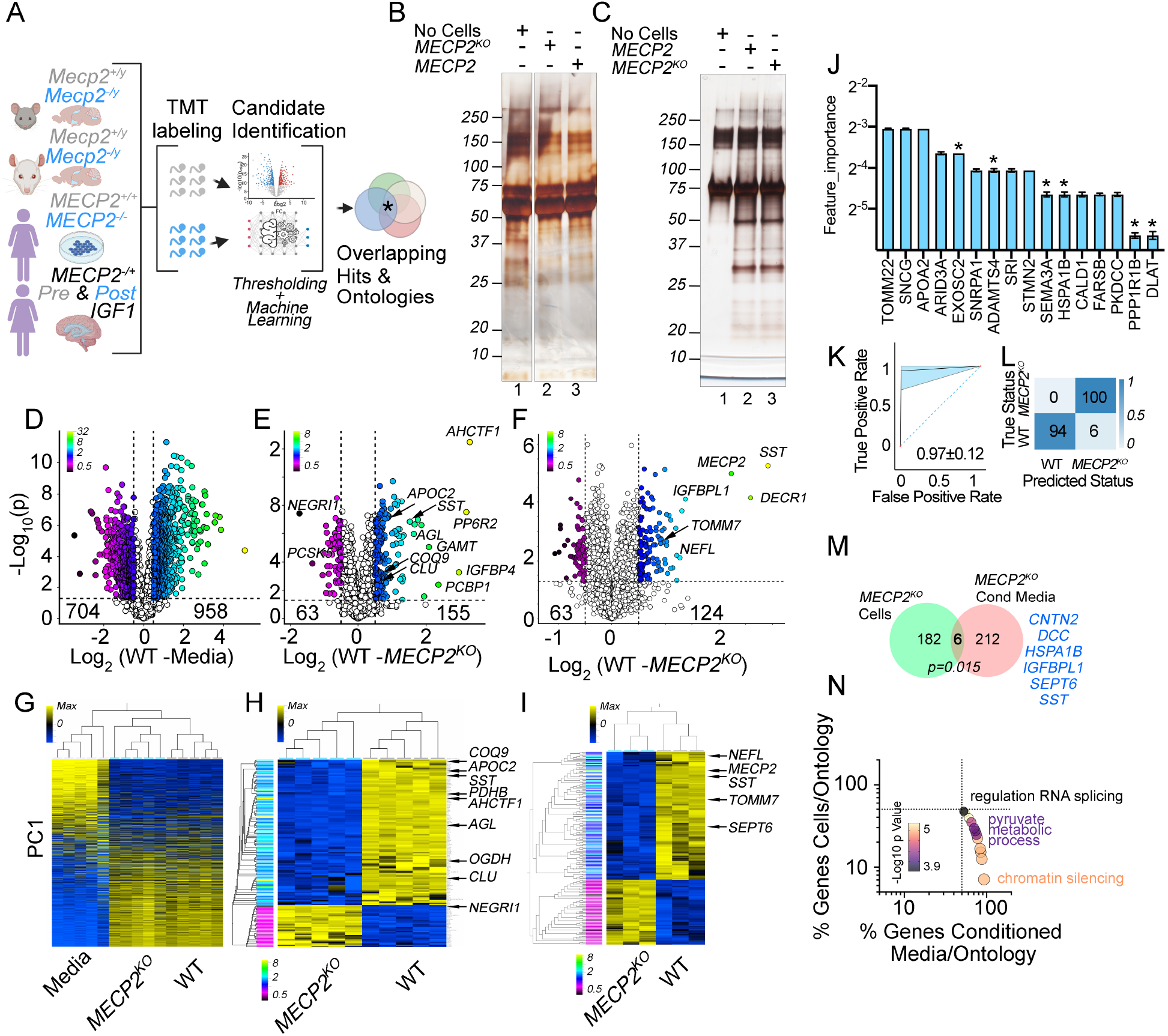
The Secreted Proteome of Post-Mitotic MECP2 Mutant Human Neurons. **A**. Experimental strategy to define a consensus Rett syndrome secreted proteome and infer conserved ontologies. **B and C**. Silver stain of two formulations of media conditioned by wild type and MECP2-null differentiated LUHMES cells, a post-mitotic human neuron line ^76^. Lane 1 in B and C represent naïve non-conditioned media. Lanes 2 and 3 depict conditioned media by wild type and mutant cells. **B** presents experiments performed with commercial N2 supplement. **C** shows experiments where the N2 supplement was custom generated from high grade purity reagents. **D** to **F**, present volcano plots of TMT mass spectrometry experiments with thresholds at log2 of 0.5-fold of change in protein abundance and a p value <0.05. Symbol color represents fold of change in linear scale (see insert). **D** presents a comparison between protein hits obtained by comparing media conditioned by wild type neurons and non-conditioned media. All hits to the right of the X axis correspond to proteins secreted by neurons (n=5). **E** shows a comparison of the wild type and *MECP2* mutant secreted proteome. All hits to the right correspond to proteins whose expression is higher in wild type than in mutant cells (n=5). **F** shows the total cellular proteome of wild type and *MECP2*-null cells used in E, n=3. **G** to **I** show clustered heat maps of hits selected in D to F. Arrows mark some cardinal hits. Rows are depicted as minimum and maximum intensities (blue-yellow scale) and annotated by log2 fold of change (rainbow scale, see table S1). **J to L** analysis of TMT data in panel E using an XGBoost machine learning algorithm. J presents main hits discriminating wild type and *MECP2* conditioned media in the decision tree. Asterisks mark proteins identified both by volcano thresholding and machine learning. K-L, performance of the machine learning protocol estimated by ROC analysis, J, and confusion matrix in I. Area under the curve in J =0.97±12. **M**, Venn diagram of the overlap between hits found in conditioned media in panel E and cellular hits in F, p value calculated with exact hypergeometric probability. **N**, shows the % overlap between the cellular proteome ontologies inferred from the datasets shown in E and F calculated with the ClueGo application (see table S1). P value estimated with exact hypergeometric probability Bonferroni corrected.

### The Secreted Proteome of a MeCP2 Deficient Neuronal Cell Line

We began using differentiated post-mitotic human neurons, LUHMES cells, where the MECP2 gene was edited by CRISPR-Cas9 ^39^. We reasoned the secreted proteome of a single cell type would define cell autonomous protein candidates for cell-type annotation of hits obtained in cerebrospinal fluids from individuals with Rett syndrome and rodent *Mecp2*-mutant models. We characterized the culture media before and after cell conditioning (Fig. 1B, compare lanes 1 with 2-3). The protein complexity of media alone prevented the identification of proteins contributed by differentiated neurons. The source of protein contaminants was commercial N2 supplements; thus, we customized a N2 supplement starting from high purity reagents. Our customized N2 allowed us to distinguish proteins contributed by either wild type or mutant neurons (Fig. 1C, compare lanes 1 with 2-3). TMT mass spectrometry of media alone identified 704 proteins (Fig. 1D and G). In contrast, cell-conditioned media revealed 958 additional proteins contributed by wild type cells (Fig. 1D and G). Next, we used this custom media formulation to compare the proteome of wild type and *MECP2* null cells. We identified 63 upregulated and 155 downregulated proteins in *MECP2* mutant cells by p value and fold-of-change volcano thresholding (Fig. 1E and H). Prominent downregulated proteins in *MECP2* mutant cells included two apolipoproteins (APOC2 and clusterin, CLU or APOJ), the nucleoporin component AHCTF1, the mitochondrial protein COQ9, as well as factors implicated in citrate cycle and glycolysis such as PDHB and OGDH ^40,41^. To these volcano selected hits, we added 10 additional proteins from a total of 16 proteins whose expression was sensitive to the *MECP2* mutation as defined by machine learning (Fig. 1J, asterisks for common volcano and machine learning hits) ^42^. Among these 10 new proteins were the mitochondrial proteins TOMM22 and the E2 component of the pyruvate dehydrogenase complex (DLAT) as well as the apolipoprotein APOA2. The performance of the machine learning algorithm was evaluated by Receiver Operating Characteristics (ROC) analysis with an area under the curve of 0.97 (Fig. 1K) and confusion matrix analysis where predicted and actual genotypic classes closely matched (Fig. 1L).

We asked whether the MECP2 secreted proteome was a reflection of the MECP2 cellular proteome. The MECP2 cellular proteome was represented by 187 proteins (Fig. 1F and I). The overlap between these two MECP2 sensitive proteomes was minor and barely significant (Fig. 1M). We found just six common hits, among them SST and IGFBPL1 (Fig. 1M). Convergence between these two datasets became evident in few significant ontological categories shared between the MECP2 secreted and cellular proteomes, one of them pyruvate metabolic process (GO:0006090 p= 3.96E-05 Bonferroni corrected, Fig. 1N). These data suggest that ontologies may be better positioned than proteomic hits to identify convergence between the secreted and cellular proteome in MECP2 gene mutations within a simple cellular system.

### Secreted Proteomes of MeCP2 Deficient Cerebrospinal Fluids in Three Species

In order to identify CSF proteomes and/or ontologies that are robust and convergent at the intra and inter-species level, we analyzed by TMT mass spectrometry the cerebrospinal fluid from wild type and *Mecp2* mutants in two rodent species and Rett syndrome individuals. We performed studies in rats (Fig. 2A-C) as well as in a large cohort of wild type and *Mecp2* null mice (Fig. 2D-F). We identified 70 and 64 CSF proteins whose expression was downregulated in mutant rat and mouse CSF, respectively (Fig. 2A, B, D, and E). These mutant CSF downregulated proteins prominently converged on subunits of high-density lipoprotein particles such as Apom, Apoa1, Apoh, and Pon1. Similarly, we identified Apoa1, Apoc1, Apoc2, and Apoe as downregulated hits in mouse *Mecp2* mutant CSF. An additional category of proteins downregulated in both species were proteins belonging to the complement and coagulation cascades (Fig. 2A, B, D, and E). To assess intraspecies robustness, we confirmed these rat CSF hits in an independent cohort of wild type and mutant rats using an orthogonal label-free mass spectrometry quantification procedure (LFQ) ^37^. This analysis identified 44 proteins whose expression was affected in mutant CSF (Fig. S2A) and confirmed the downregulation of apolipoproteins and complement factors in *Mecp2*-null CSF (Fig. S2A-C, Apoa4, Apob, Apoc3, Pon1, and C9). Rodent apolipoprotein and complement cascade hits could not be attributed to blood contamination of the CSF as evidenced by albumin, immunoglobulins, or hemoglobin species, which failed to co-cluster with apolipoproteins and complement (Fig. 2B and E, S2A-B, Table S2). Synaptic proteins were prominent among factors upregulated in the mouse Mecp2 mutant CSF (Fig. 2D-E). These synaptic proteins include, but were not limited to, Snap25, Stx1b, Stxbp1, Syn1, and Syn2 (Fig. 2D-E) ^43^. Finally, we also identified mitochondria proteins within both the significant up- and downregulated proteomes in mutant rat and mouse CSF (Fig. 2D-E).

**Figure 2.**
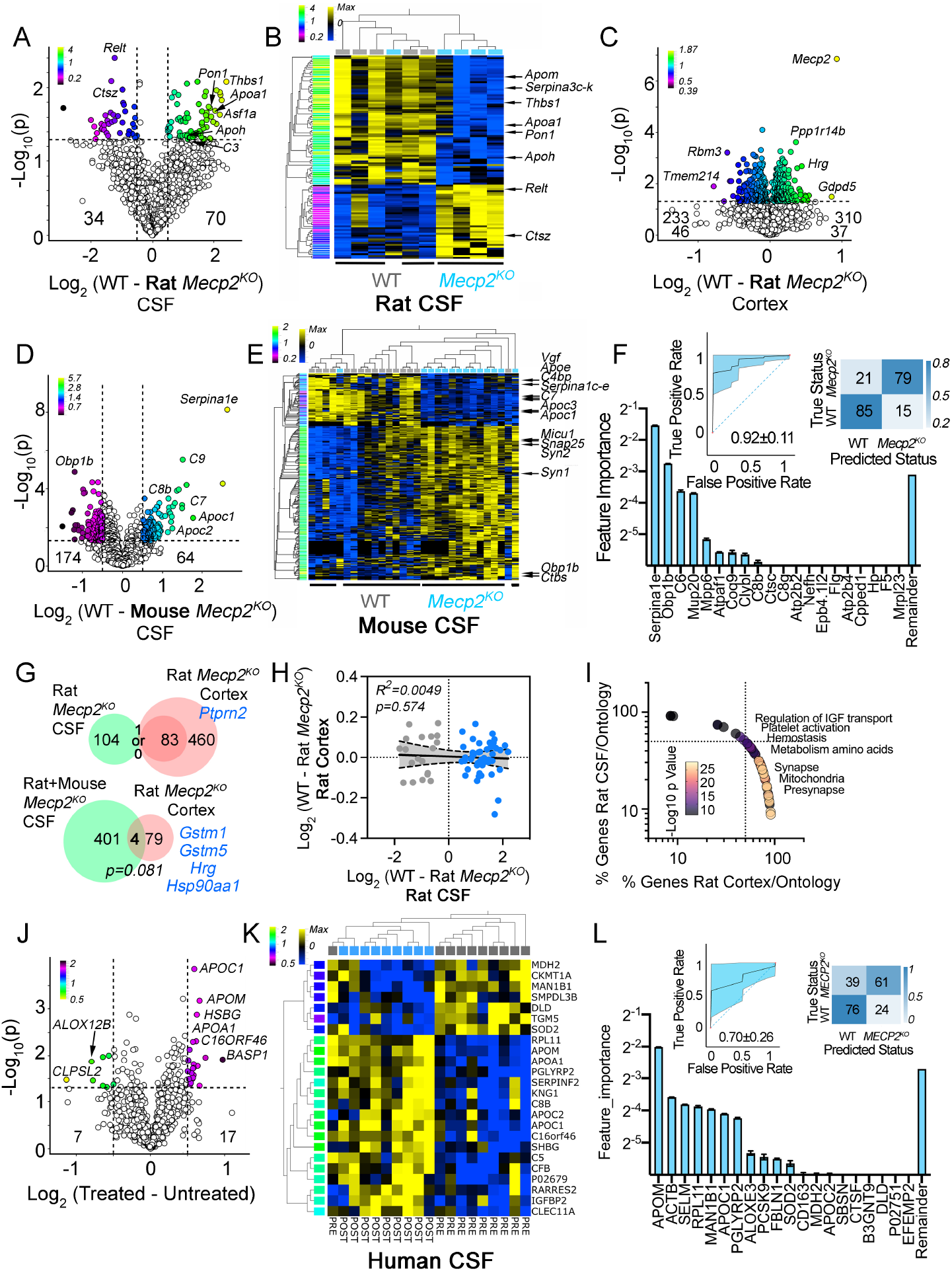
The Rett Syndrome CSF Proteome Across Three Species. **A** volcano plot of TMT mass spectrometry determinations in rat cerebrospinal fluid. Cutoffs at log2 of 0.5 fold of change in protein abundance and a p value <0.05 n=5. **B** shows clustered heat maps of hits selected in A. For A and B, see legend to Figure 1 for additional details. **C** depicts volcano plot of rat cortices analyzed by TMT mass spectrometry. Shown are hits selected by p value <0.05 see table S2 for hit lists at p<0.01. n=5. **D** and **E** show mouse CSF TMT volcano plot and heat map of selected hits at cutoffs of log2 0.5 fold of change in protein abundance and a p<0.05. n=16 wild type and 14 *Mecp2*-null mice. **F** analysis of TMT data presented in panel D using an XGBoost machine learning algorithm. Main hits discriminating wild type and *Mecp2* mutant CSF proteomes in decision tree are shown. Inserts show performance of the machine learning protocol estimated by ROC analysis and confusion matrix. Area under the curve in J =0.92±11. **G**, Top Venn diagram shows the overlap between *Mecp2*-sensitive rat CSF and rat cortex hits using two thresholding criteria p<0.05 and p<0.01. Bottom Venn diagram compares *Mecp2*-sensitive hits in rat and mouse CSF pooled together with *Mecp2*-sensitive rat cortex hits. **H** depict the correlation in expression of *Mecp2*-sensitive hits in rat CSF and rat cortex. **I**, shows the % overlap between the cellular proteome ontologies inferred from the datasets shown in A and C calculated with the ClueGo application (see table S2). **J** and **K** show Rett syndrome female individual CSF TMT volcano plot and heat map of selected hits at log2 of 0.5 fold of change in protein abundance and a p<0.05 comparing before and after IGF-1 treatment. n=10 before treatment and 9 after treatment. **L** analysis of TMT data presented in panel J using an AdaBoost machine learning algorithm. Main hits discriminating CSF before and after treatment in decision tree are shown. Inserts show performance of the machine learning protocol estimated by ROC analysis and confusion matrix. Area under the curve in insert ROC analysis 0.70±0.26.

We further scrutinized mouse and rat CSF datasets using machine learning algorithms. We sought to identify additional proteins sensitive to *Mecp2* deficiency that could otherwise escape detection by volcano thresholding. In addition, we reasoned proteins categorized as apolipoproteins, synaptic, complement-related, or mitochondria-annotated should emerge as priority hits in non-linear decision trees segregating wild type and Mecp2 mutant CSFs. Both mouse and rat CSF machine learning analyses identified complement factors, mitochondrial proteins (Atpaf1, Clybl, Coq9, and Mrpl23), as well as synaptic proteins as priority hits (Actr2, Atp2b2 and Nefh, Fig. 2F and S2C-E). We validated the performance of machine learning approaches asking their capacity to identify Mecp2 as a priority protein hit in a proteome dataset of wild type and Mecp2 mutant rat cortex (Fig. S2D). We identified Myg1, a mitochondrial protein ^44^, and Mecp2 as the top-two most important classifiers to discern between wild type and *Mecp2* mutant brain tissue (Fig. S2D). All these machine learning analyses performed satisfactorily as determined by area under the curve in ROC analysis (0.72-0.92, Fig. 2F, and S2) and/or confusion matrices (Fig. 2F, and S2). Thus, interrogation of CSF proteome datasets with boosting mathematical algorithms provides similar answers as to protein families enriched in rat and mouse mutant CSFs.

The above-described changes to the *Mecp2* mutant CSF could closely parallel brain proteome modifications. Alternatively, changes to the CSF proteome could be in proteins different from those in the Mecp2 brain proteome yet both *Mecp2* proteomes representing alterations in the same compartment or pathway, a converging ontology. To address this question, we compared the rat *Mecp2* mutant cortex proteome of animals where we simultaneously collected CSF. Volcano thresholding by p value and fold of change identified 83 protein whose expression was modified in *Mecp2* mutant cortex among 6752 proteins quantified by TMT (Fig. 2C). None of the *Mecp2*-sensitive cortical proteins overlapped with rat CSF hits (Fig. 2G). We reasoned that a lack of overlap could emerge from a *Mecp2*-sensitive cortex proteome with small expression differences as compared to CSF due to the complexity of the former. Thus, we relaxed thresholding to include proteins whose expression was significantly different (p<0.05) without applying a fold-of-change cutoff. This resulted in a relaxed dataset of 543 cortical proteins in whose expression may be modified in *Mecp2* mutants (Fig. 2C). However, this enlarged and relaxed cortical dataset produced only one overlapping hit with CSF (Fig. 2G, Ptprn2). An alternative model to account for the lack of overlap between *Mecp2* cortex and CSF proteomes could come from sparse hits identified in mutant rat CSF. Yet comparison of a 405 proteins pooled rat and mouse *Mecp2*-sensitive CSF proteome, obtained by thresholding and machine learning, yielded a non-significant overlap of just four hits (Fig. 2G). We expanded our analysis searching for CSF hits in *Mecp2* mutant rats that were also quantified in rat cortices. We found 67 proteins shared between the rat cortical dataset and the rat CSF *Mecp2*-sensitive proteome. These 67 proteins showed no correlation in their expression levels (Fig. 2H). Finally, we evaluated whether the *Mecp2*-sensitive CSF and cortical proteomes could overlap at the ontology level. To evaluate this hypothesis, we compared whether the rat *Mecp2*-sensitive cortex and CSF proteomes could converge on similar ontologies. We identified significant ontological categories only if the *Mecp2*-sensitive CSF proteome was analyzed in conjunction with the relaxed *Mecp2* cortical proteome of 543 proteins. Under these conditions, we found overlapping ontologies including the synapse, which was represented by 75% of the annotated proteins contributed by the cortical and 25% by the CSF Mecp2-sensitive proteomes (Fig. 2I, GO:0045202, Bonferroni corrected group p=2.467E-22). These results argue that ontologies predicting putative brain disease mechanisms can be inferred from changes in the CSF proteome. However, there is a limited capacity to predict specific protein candidates in *Mecp2* mutant CSF from brain proteomes and vice versa.

The *Mecp2* mutant rodent CSF differs from the human Rett CSF in that the former represents brain tissue homogenously deficient in Mecp2 protein. In contrast, the human CSF proteome reports a genetically mosaic female brain ^3-5^. We studied a cohort of individuals with Rett syndrome where CSF was collected as part of a phase I clinical trial ^45^. These participants were subjected to extended treatment with recombinant IGF-1 during the trial. CSFs were collected from 12 individuals. In nine participants, fluids were collected before and after treatment; in two participants, collections occurred only before treatment; and in three participants, CSF was sampled only after IGF-1 treatment. The phase I clinical trial did not include typically developing control participants due to ethical constraints ^45^. Even though IGF-1 treatment did not improve clinical outcomes in Rett subjects ^35^, we reasoned that if IGF-1 treatment were to modify some aspects of CSF proteome, it should do so by changing CSF proteins whose expression was *Mecp2*-sensitive in rodents. In addition, we hypothesized that any IGF-1-induced proteome modifications in individuals with Rett syndrome should be in the opposite direction of what we observed in *Mecp2*-null rodents and conditioned media from *MECP2* null neurons. The Rett CSF proteome revealed a discrete number of proteins sensitive to IGF-1 (Fig. 2J-L). We identified by volcano thresholding 7 proteins whose expression was decreased after IGF-1 treatment and 17 proteins whose expression was increased (Fig. 2J-K). Importantly, the most prominent among the IGF1-upregulated proteins were the high-density lipoprotein proteins APOA1, APOC1 and APOM; a change precisely in the opposite direction of what we found in mouse and rat *Mecp2* mutants. Apolipoproteins APOC1, APOC2, and APOM as well as the apolipoprotein regulatory factor PCSK9 were also identified by machine learning (Fig. 2L). In fact, APOM was assigned the top priority as a discriminatory factor in a decision tree segregating Rett participants by their IGF-1 treatment (Fig. 2L). Our data suggest that discrete CSF proteome changes correlate with IGF-1 treatment in Rett female individuals. In the case of apolipoproteins, protein identity and the direction of change in humans can be informed from the study of *Mecp2* mutant CSF proteomes in preclinical animal models.

### Composite Ontologies of *Mecp2* Mutant Cerebrospinal Fluids and Conditioned Media

The secreted proteomes from human cultured neurons, rat, mouse, and human mutant CSFs revealed common proteins across these diverse experimental systems. These proteins belong to apolipoproteins, complement, or mitochondrial pathways (Fig. 3A). We asked if these *Mecp2*-sensitive hits overlapped just as isolated occurrences or, instead, secreted proteomes obtained from each experimental system sampled a common ontological space. We tested this hypothesis using ClueGO, an application that performs composite and comparative enrichment tests based on hypergeometric distributions ^46^. To test the robustness of our ontological predictions, we used HumanBase, a genomics data-driven Bayesian machine learning algorithm that identifies functional modules in tissues and cells ^47^. We collated proteins from each of the four *Mecp2*- and *MECP2*-deficient experimental systems, selected by volcano thresholding plus machine learning (Fig. 1A), to identify a space of shared ontologies (Fig. 3A). We simultaneous queried an ontological database composite with ClueGo (GO CC, REACTOME, KEGG and WikiPathways). Each of the four mutant proteome datasets was tagged in ClueGo to discern their individual contributions to each ontology. We identified a space of 87 ontologies significantly represented in all the *Mecp2*- and *MECP2*-sensitive proteome datasets (Bonferroni corrected p values <10E-3, Fig. 3B). These ontologies revealed a significant enrichment of hits in mitochondrial compartments and pathways, pyruvate and aminoacid metabolism, complement subunits, HDL lipoproteins, and synapse-related ontologies (Fig. 3B). Importantly, these ontologies were qualified as non-dataset specific, as each mutant proteome dataset contributed less than 50% of hits to each one of these ontologies (Fig. 3C-D). For example, the HDL particle ontology was made up by 34, 26, 16 and 25% of hits derived from mouse, rat, neuron conditioned media, and human Rett CSF, respectively (Fig. 3D, GO:0034364, Bonferroni corrected group p value=2.81E-13). We confirmed these ontological findings using HumanBase, where we identified lipid and cholesterol transport ontologies as the most significantly enriched functional modules (Fig. 3E). This outcome was similar if we performed HumanBase analyses either with astrocyte- or neuron-centric queries (Fig. 3E). Finally, we confirmed that *Mecp2*- and *MECP2*-sensitive proteomes were enriched in HDL, synaptic, and mitochondria annotated proteins interrogating different databases. We used the curated HDL proteomes database, the SynGo knowledge base of annotated synaptic proteins, and the Mitocarta 3.0 database of annotated mitochondrial proteins (Fig. 3F-G) ^41,43,48^. The collated *Mecp2*- and *MECP2*-sensitive proteome contained 80 proteins in common with the HDL proteome (Fig. 3F). This represents a significant 7.9-fold enrichment above what is expected by chance (Fig. 3F, p<9.26E-49). Among these overlapping HDL proteins, we found diverse apolipoproteins, complement subunits, antiproteases of the serpin family, and factors such as clusterin (Clu), and PCSK9. All these overlapping HDL components formed an interconnected network of protein-protein interactions as determined with the Genemania application (Fig. 3F) ^49^. Similarly, the collated *Mecp2*- and *MECP2*-sensitive proteome was also significantly either enriched in interconnected synaptic or mitochondrial proteins >2-fold above what is predicted by chance. These findings demonstrate that Rett syndrome CSF proteomes from diverse species converge on a common set of ontologies. These ontological findings demonstrate a varied and novel set of molecular phenotypes associated to Rett syndrome. These Rett syndrome molecular phenotypes offer a distinctive path for disease biomarkers.

**Figure 3.**
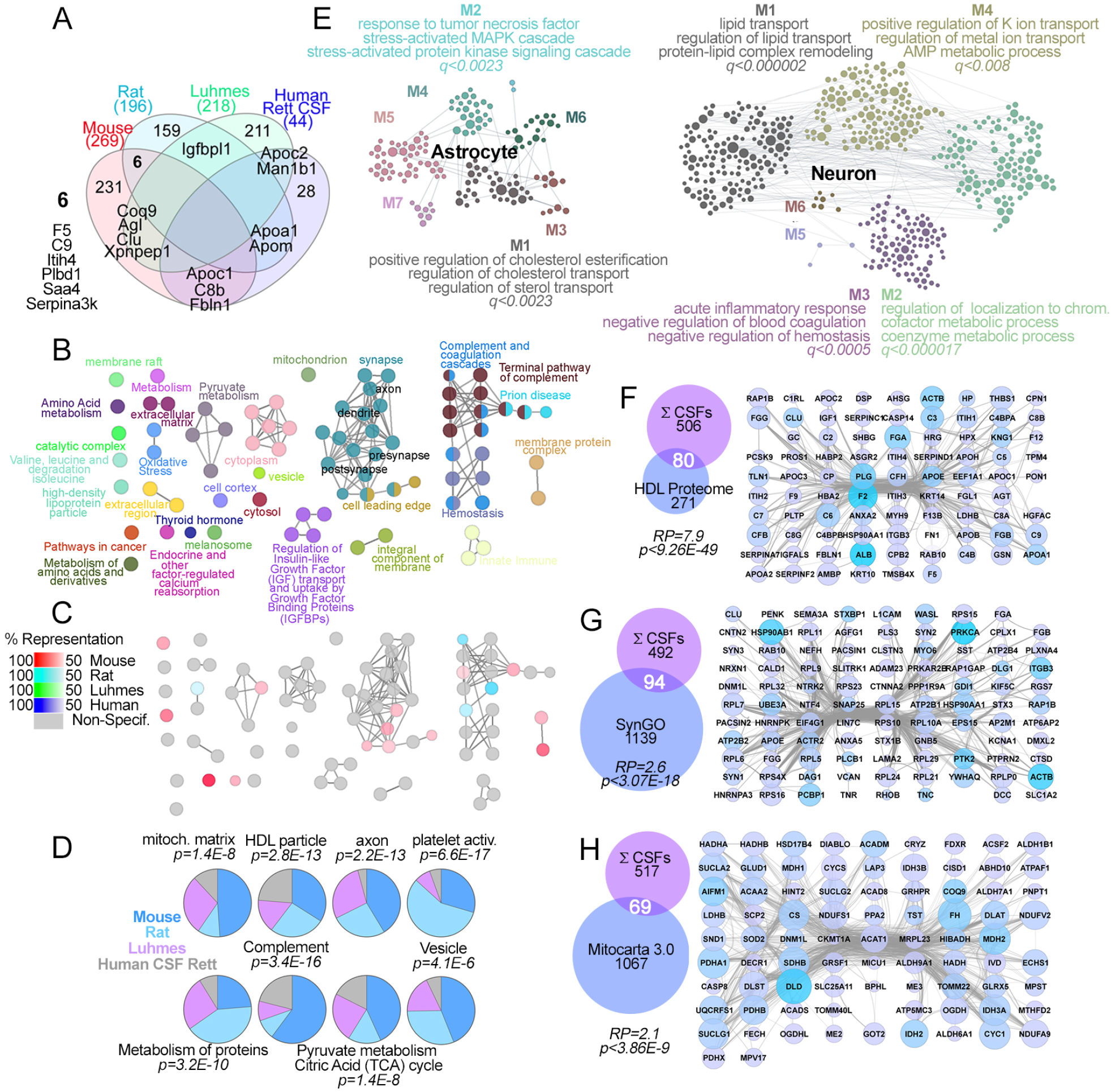
Convergent Ontologies Inferred from Rett Syndrome CSF and CSF-mimic Proteomes. **A**, Venn diagram depicted hit overlaps among the four experimental systems studied. Hits represent the sum of volcano thresholding- and machine learning-selected hits. **B and C**. Integrated gene ontology analysis of the four datasets in A annotated with the experimental system that originated the dataset. Nodes represent individual ontologies. GO_CellularComponent, KEGG, Reactome, and WikiPathways were queried with the ClueGo application. All ontologies have p<0.001. Exact hypergeometric probability Bonferroni corrected. **C**, shows the percent of contribution of each experimental system to each ontology. Grey denotes ontologies represented by all four experimental systems. **D**, pie charts of the percent contribution of each experimental system to the top ontologies identified in B-C, p values hypergeometric probability Bonferroni corrected for the ontology. **E**, shows ontology analysis using the same datasets in B-C but using the Bayesian engine HumanBase. Analyses were performed either with astrocyte- or neuron-centric queries. Nodes correspond to genes and their functional interrelationships. Nodes are grouped into clusters M(n), prioritized by q value calculated using one-sided Fisher’s exact tests and Benjamini–Hochberg corrections. Most significant cluster is M1. **F-H**, show Venn diagrams and protein-protein interaction networks of the sum of datasets presented in A overlapping with the curated HDL proteome in F, the Synapse knowledge database of annotated genes in G, and the Mitocarta 3.0 annotated mitochondrial proteins in H. Venn diagram p values calculated with exact hypergeometric probability and the representation factor (RP) estimates overlap beyond what is expected by chance. See Table S3 for all ontological data.

### Ontologies Inform Putative Biomarkers of Rett Syndrome

We used convergent Rett molecular ontologies to inform a selection of proteins for confirmatory studies and to assess their potential as disease biomarkers. The HDL lipoprotein proteome was the most significantly enriched ontology among all the mutant secreted proteomes (Fig. 3B, E and F). We performed absolute quantification (AQUA) of proteins by mass spectrometry to confirm expression changes in HDL apolipoproteins in independent cohorts of rat (Fig. 4A) and mouse CSF from wild type and mutant animals (Fig. 4B-C) ^50^. We also confirmed CSF hits in Rett syndrome individuals before and after IGF1 treatment by electrochemiluminescent ELISA assays (Fig. 4C) ^51^.

**Figure 4.**
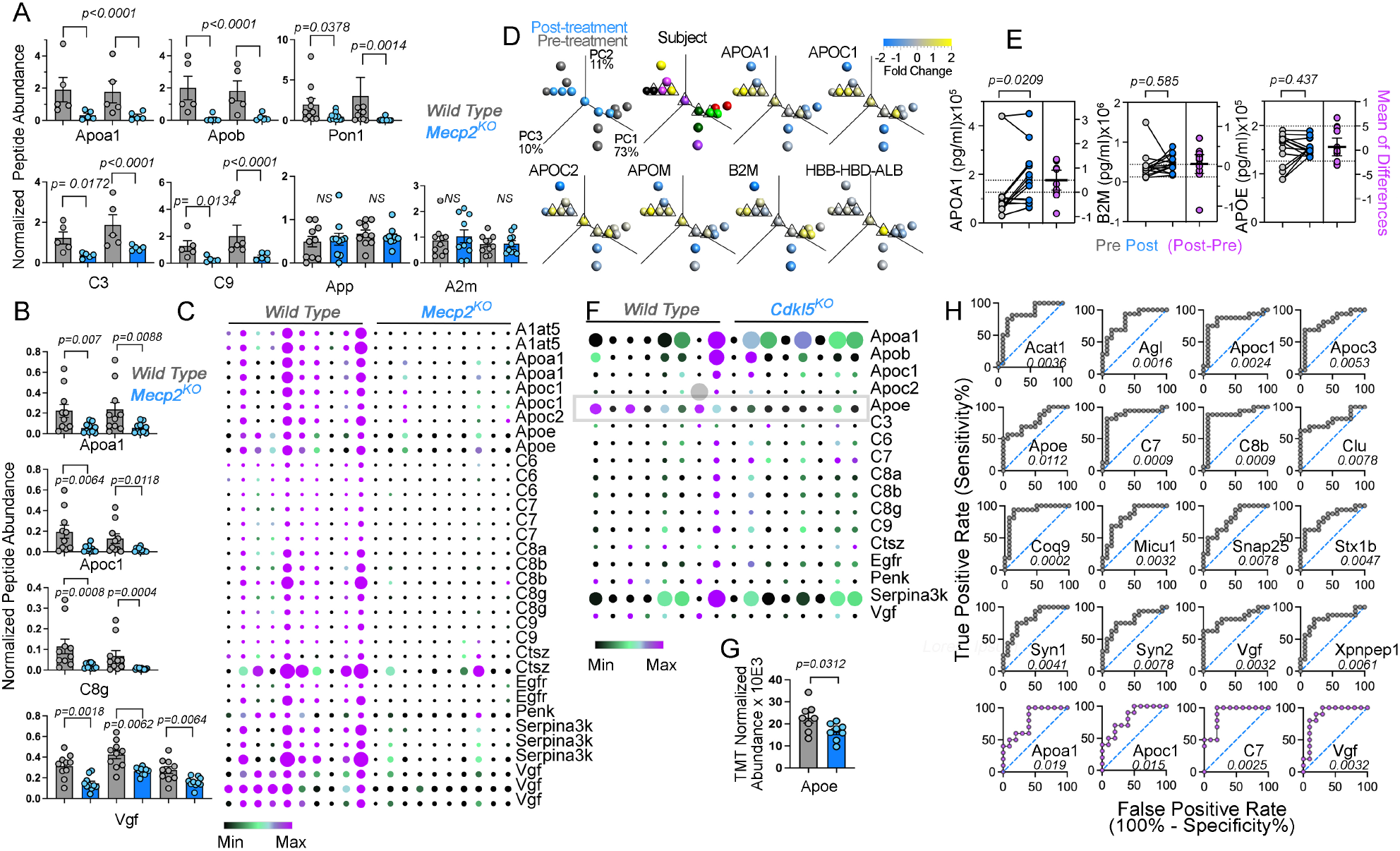
Ontologically Selected and Confirmed Rett Syndrome CSF Proteome Hits Perform as Putative Biomarkers of Disease. **A** and **B** shown independent confirmatory analyses using isotopologue peptides mapping the primary sequence of the indicated proteins using AQUA mass spectrometry in rat CSF (A) or a modified AQUA approach in mouse CSF (B). In **A**, femtomoles of the endogenous CSF peptide were normalized to a randomly chosen control sample. In **B**, the ratio of the CSF endogenous peptide to the isotopologue peptide were used to quantify relative analyte abundance. Gray bars correspond to wild type CSF and blue bars *Mecp2* null CSF. All analytes were measured independently with 2-3 isotopologue peptides as standards. In A two batches of 5 rats of each genotype were used while in B, one batch of 10 mice of each genotype were analyzed. **C** depicts a heat map of all the modified AQUA determinations performed in mouse CSF samples that showed significant differences between genotypes. Every isotopologue peptide correspond to a row. Data are depicted as row median divided by the row median absolute deviation both as heat map and by symbol size. **D**, principal component analysis of Rett syndrome participants before and after treatment (gray and blue symbols, respectively), every subject is color coded and the after-treatment PCI coordinates are indicated by a callout triangle. Analyte expression levels from a TMT mass spectrometry quantification were mapped to the PCI coordinates as a heat map. Apoliproteins increased expression after IGF-1 treatment. Note controls do not experience changes (B2M, HBB, HBD, ALB). **E**, MesoScale ELISA confirmation of APOA1 levels in Rett individuals CSF before and after IGF-1 treatment. **F** and **G**, a heat map of ontology selected and confirmed analytes in B-C tested in the CSF of *Cdkl5*-null mice. Data correspond to the normalized abundance measured by TMT mass spectrometry. **G** shows APOE, the only analyte whose expression was modified in *Cdkl5*-null CSF, shown by the gray box in F. **H**, ROC analysis of selected mouse CSF *Mepc2*-sensitive hits either quantified by TMT mass spectrometry (grey symbols) or by modified AQUA (purple symbols). Number represents p value that tests the null hypothesis that the area under the curve=0.50 (non-discriminatory). P values in A, B, E and G were calculated with a two-sided permutation t-test. See table S4 for data to panels B-C and F-G.

AQUA quantification of the HDL lipoprotein particle components Apoa1, Apob, Pon1, C3, and C9 revealed decreased levels in mutant as compared to wild type rat CSF (Fig. 4A). Levels of loading controls, App and A2m, were not affected by genotype in rats (Fig. 4A). These findings were extended to Rett mouse models (Fig. 4B-C). We used isotopolog peptide standards to measure proteins associated to HDL particles in mouse CSF. Diverse apoliproteins (Apoa1, Apoc1, Apoc2, Apoe), complement subunits (C6, C7, C8a, C8b, C8g, C9), and antiproteases (A1at5, Serpina3k) were reduced in *Mecp2* deficient animals compared to controls (Fig. 4B-C). Similarly, the levels of other secreted proteins such as the neurosecretory protein VGF, proenkephalin-A (Penk), cathepsin Z (Ctsz), or the transmembrane epidermal growth factor receptor (Egfr) were robustly and significantly reduced in the CSF of mutant mice (Fig. 4C). These results independently validate our TMT findings in the CSF of two rodent models of Rett syndrome.

We next focused on the nine participants where CSF samples were obtained before and after IGF1 treatment. We deployed principal component analysis to assess individual and group responses to IGF1. The main variable segregating participant CSF proteomes was their identity rather than treatment itself (Fig. 4D). Second, treatment modified a participants’s CSF composition discretely, mostly driven by changes in the levels of apolipoproteins (Fig. 4B). Importantly, the levels of blood proteins such as hemoglobins (HBB and HBD), albumin (ALB), and immunoglobulins could not account for the changes in apolipoprotein protein levels (Fig. 4B). We confirmed that IGF1 treatment increased the content of APOA1 in Rett participant’s CSF by ELISA (Fig. 4C). In contrast, two proteins whose expression was not modified by treatment in TMT mass spectrometry quantifications, APOE and B2M, did not change their levels after treatment in ELISA assays (Fig. 4C). These results confirm protein hits and ontologies in human and rodent Rett syndrome models.

To evaluate the potential of ontologically selected *Mecp2*-sensitive hits to serve as disease biomarkers, we addressed the following questions. First, do CSF *Mecp2* hits discriminate between genetic forms of autism spectrum disorder? We selected a null mutation in the *Cdkl5* gene, which is causative of the CDKL5 deficiency disorder, an X-linked neurodevelopmental disorder. The behavioral and brain anatomy phenotypes of a mouse model of this syndrome closely mimic those in Rett syndrome mouse models ^33^ (Fig. 4F-G). We found that the CSF proteome of *Cdkl5* mutants was different from *Mecp2* mutant CSF, as indicated by the 16 different proteins selected from Mecp2 CSF whose levels remained unchanged in *Cdkl5* null CSF. The only exception was Apoe, which was decreased in both *Mecp2* and *Cdkl5* mutant CSF (Fig. 4G). These findings demonstrate that *Mecp2* CSF hits discriminate between phenotypically related forms of neurodevelopmental disorders. Second, we interrogated whether selected *Mecp2* CSF proteins were expressed in disease-relevant cell types (Fig. S3A-B). HDL apolipoproteins such as Apoa1, Apoe, Pon1, and Clu were expressed in neurons and astrocytes (Fig. 4A). Single cell RNAseq data showed that transcripts encoding these HDL proteins were expressed in diverse populations of glutamatergic and GABAergic neurons across the multiple layers of the cortex and hippocampus (Fig. S3B). Apoc1 mRNA or its protein were undetectable in neurons and glia but present in plasma (compare Fig. S3A and B). In contrast, complement components (C7, C8b, C8g, and C9), growth factors (Igf1 and Vgf), mitochondrial proteins (Acat1, Coq9, and Micu1), and synaptic annotated proteins (Snap25, Stx1b, Syn1, and Syn2) were expressed in diverse glial and neuronal cell populations to a different degree (Fig. S3B). For example, C8b and C9 mRNAs were expressed in a discrete neuronal population whereas C7 was broadly expressed in cortex and hippocampus (Fig. S3B). Thus, ontologically selected protein hits are expressed in disease-relevant neuronal and glial cell types. Finally, we determined if selected proteins could distinguish subjects by genotype. We performed ROC analysis focusing on the *Mecp2* mutant mouse CSF hits as the animal cohort of the biggest size, the mouse TMT dataset (Fig. 4H). Each ontologically selected analyte efficiently distinguished genotypes irrespective of if they belonged to either the HDL lipoprotein, synapse, or mitochondrion category. In fact, all analytes ROC area under the curve were between 0.77 to 0.9 with significant p values (Fig. 4H). In other words, these analytes have a 77 to 90% chance to distinguish wild type and mutant CSF. The ROC performance of an analyte was similar whether TMT or AQUA datasets were analyzed (Fig. 4H, compare grey and purple symbols). None of these validated analytes experienced modifications in their mRNA expression across diverse brain regions in *Mecp2* mutant mice, indicating that the utility of these analytes as putative biomarkers of *Mecp2* gene defect is restricted to their protein levels in CSF (Fig. S4). These findings demonstrate that CSF analytes identified in rodent models of Rett syndrome are sensitive, conserved, and represent specific candidates for Rett biomarkers with potential for human applications.

## Discussion

Here we demonstrate that a X-linked neurodevelopmental disorder, Rett syndrome, reproducibly and distinctively impacts the composition of the CSF. We defined a consensus proteome and ontological categories shared by four experimental systems across three species deficient in *Mecp2/MECP2*. These proteomes converged on proteins annotated to HDL lipoproteins, complement cascade, mitochondrial compartments, citrate cycle/pyruvate metabolism, as well as synapse compartments. The robustness of our findings is founded on the multipronged nature of our experimental design, which includes diversified in vitro and in vivo systems, multiple species studied, distinct mathematical processing of datasets capturing similarly annotated proteins, replication across different proteomic platforms (LFQ and TMT), and replicability across two sites for CSF collection and mass spectrometry analysis (Fig. S1). Although, the different mutant CSF proteomes produced discrete overlap across individual analytes, they all shared significant overlap at the ontology level. We used these convergent ontologies to inform the selection of analytes for orthogonal confirmatory efforts. These confirmatory approaches independently validated our LFQ and TMT findings. Confirmed analytes provided a proof of principle to the use of convergent ontologies as a strategy to select analytes predicted to report a mutant genotype across species and quantification platforms. For example, even though Apoa1 was not identified as a significant hit in the mouse CSF TMT proteome, selection of Apoa1, based on the ontology to which it belongs, predicted and resulted in robust confirmation across all species studied and platforms used. Ontologically selected hits performed well as putative biomarkers as determined by ROC analysis and the capacity of multiple *Mecp2*-sensitive hits to discriminate *Mecp2* mutant CSF from another phenotypically related neurodevelopmental disorder, the CDKL5 deficiency disorder. We propose that *Mecp2* mutant CSF ontologies inform robust CSF analytes to act as Rett syndrome biomarkers in humans.

The mechanisms that account for the changes in the secreted proteome described here have not been explored yet. However, we reasoned that if the *Mecp2* secreted proteome were to be caused directly by a *Mecp2*-dependent transcriptional defect, there should be parallel modifications in the cellular and secreted proteomes. We found that this is not the case. The CSF and brain cortex proteomes did not correlate. In fact, transcriptomic analysis of several ontologically selected CSF protein hits showed that none of these proteins exhibited correlated modifications in their mRNA levels in brain. Similarly, the neuronal cell and conditioned media proteomes poorly overlapped. These results argue that indirect mechanisms downstream of *Mecp2*-dependent transcription, such as network activity, likely drive the secreted proteome phenotypes.

We have minimized the possibility that accidental plasma contamination of the CSF is a driving factor for some of the CSF expression differences observed. However, CSF protein composition is defined by factors that normally transcytose from the plasma to the CSF plus contributions from neuronal and non-neuronal cells in the brain parenchyma, the choroid plexus, and ependymal cells ^52,53^. Therefore, our findings likely represent contributions of diverse cell types in brain to the *Mecp2*-sensitive CSF proteome. With few exceptions, many of our CSF *Mecp2*-sensitive proteins could be attributed to multiple cell types. For example, Apoe and Clu (Apoj) could be ascribed to secretions from astrocytes or the choroid plexus, where Apoe and Clu (Apoj) rank among the most expressed mRNAs; they could be ascribed to neurons, where these mRNAs are also expressed yet at lower levels. Importantly, Apoe brain levels are locally controlled without contributions from plasma ^54^. We directly tested the hypothesis that the expression of apolipoproteins is cell-autonomously controlled in neurons as demonstrated by the reduced levels of Clu (Apoj) in the conditioned media of human postmitotic neurons. On the other extreme, Vgf and Igf1 mRNAs are expressed in neurons with preferences for neuronal cell types. Such is the case of Igf1, which is mostly expressed in GABAergic interneurons and is minimally or not expressed in glia, endothelial cells, and the choroid plexus ^55,56^. Thus, the *Mecp2* secreted proteome offers multiple analytes to assess phenotypes in multiple brain cell types lacking *Mecp2*.

All *Mecp2* secreted proteomes converged on robust ontologies. Proteins annotated most significantly to HDL lipoprotein, complement, synapse, mitochondria, and mitochondrial pathways such as citrate cycle/pyruvate metabolism ontologies. These consensus ontologies likely point to pathogenic mechanisms in Rett syndrome. For example, the effects of *Mecp2* mutations on synaptic morphology, function, and plasticity have been extensively documented ^4,57^. However, HDL and mitochondrial ontologies have received less attention. HDL particles are assembled by astrocytes and microglia. These lipoproteins transport cholesterol between glial cells and neurons. Thus, a possible mechanism to account for the decreased levels of HDL apolipoproteins in CSF is either a decreased production/secretion by *Mecp2* deficient glial cells or an increased clearance by cells that express HDL receptors in brain, such as neurons ^58,59^. We favor the decreased HDL production model as it can explain the observed increased cholesterol content in brain at postnatal day 56, despite decreased expression of cholesterol synthesis enzymes and decreased *de novo* cholesterol synthesis ^60^. We postulate that decreased HDL lipoproteins levels in CSF may be a factor contributing to the accumulation of cholesterol in brain and the concurrent inhibition by product of cholesterol synthesis. A second ontology strongly represented in our datasets is mitochondria compartments and pathways. Pyruvate and lactate are increased in Rett syndrome individual’ CSF, and Krebs cycle metabolites are increased in the brain of *Mecp2* mutant mice. This suggests connections between CSF glycolysis and Krebs cycle ontologies and proteins and these metabolites ^29,30^. However, there are not enough studies to tie together our observations in CSF with potential models of mitochondrial dysfunction in *Mecp2* mutant cells ^61-65^. Our findings support the idea that *Mecp2* mutant CSF ontologies predict putative brain mechanisms disrupted by mutations in *Mecp2*. We propose that Rett syndrome is a synaptic and metabolic disorder of neurodevelopment.

## Materials and Methods

### Rat and Sample Collection

All experiments were carried out in accordance with the European Communities Council Directive (86/609/EEC) and with the terms of a project license under the UK Scientific Procedures Act (1986). The *Mecp2*^*-/y*^ rats were maintained by crossing *Mecp2*^*+/-*^ females with wild type Sprague Dawley males. Animals were maintained on 12-hour light/dark cycles with free access to normal rat food. WT and *Mecp2*^*-/y*^ rats at 25 days of postnatal age were weighed and assessed for the development of the RTT-like phenotypes prior to surgery. Rats were anaesthetized using intraperitoneal administration of an injectable cocktail of medetomidine (0.5mg/kg) and ketamine (75mg/kg). Once the animal was deeply anaesthetized, as indicated by the absence of withdrawal reflexes (tail and limbs) and the eye positioning reflex, the surgical area was shaved and the animal was secured in the stereotaxic frame with the head tilted at roughly 45º. The surgical area was then cleaned with Hibiscrub and a surgical drape was placed around the operating area with a hole to expose only the surgical area. A skin incision along the midline of the skull extending from between the eyes to 3-4 cm caudally to make sure the back of the neck is fully exposed. The fascia and the superficial and deep layers of the neck muscles were then dissected to expose the membrane of the dura mater at the atlanto-occipital joint between the occipital condyles and the rostral facets of atlas. The cisterna magna was then carefully pierced by a pulled glass pipette (1 cm long) connected to a 2.5ml syringe through 30 cm of PE-50 tubing. A small volume of CSF entered the glass pipette through the capillary action and the flow was maintained by gently pulling the plunger. The CSF was collected into cryoprotective tubes and snap-frozen immediately in liquid nitrogen. Animals were then given a lethal dose of anesthesia, decapitated and the brain was exposed and the areas of interest were dissected and snap-frozen in liquid nitrogen.

### Mice and CSF Collection

Animal husbandry and euthanasia was carried out as approved by the Emory University Institutional Animal Care and Use Committees. C57BL/6J male mice (The Jackson Laboratory #000664). Mecp2, *Mecp2*^*tm1*.*1Bird*^, and Cdkl5-deficient mice, *Cdkl5*^*tm1*.*1Joez*^, were obtained from the The Jackson Laboratory stocks #003890 and 021967, respectively. All animals were of 6 weeks of age. Animals were maintained on 12-hour light/dark cycles with free access to mouse chow.

Our terminal CSF collection method was adapted from a previously published protocol ^66^. Mice were deeply anaesthetized by intraperitoneal injection of a mixture of ketamine (73.5 mg/kg; Akron, USA), xylazine (9.2 mg/kg; Bayer Pharma, Germany), and acepromazine maleate (2.75 mg/kg; Boehringer Ingelheim, USA) in 0.9% (v/w) NaCl. The back of the neck overlying the occiput were first shaved then cleaned and disinfected with 70% ethanol. Using the thumb and index finger, the mouse was placed prone with the neck in flexion on a 15 mL conical tube at approximately 45-degree angle to access the cisterna magna using landmarks between occipital protuberances and the spine of the atlas. A Hamilton syringe containing 30 G needle was inserted through the skin at a 45-degree angle with the horizontal, to reach a depth of approximately 4 mm into the cisterna magna for CSF collection without need for an incision. The syringe was kept stable without any lateral movement and 4-12 µl or clear CSF was drawn into the syringe by slow and smooth aspiration. The CSF was immediately spun down for 30 seconds and clear CSF was inspected with the naked eye and frozen immediately on dry ice. Frankly blood contaminated samples discarded.

### Human Subjects

Clinical features of the cohort used in these studies are described by Khwaja et al.^45^. The referred study was approved by the Institutional Review Board of Boston Children’s Hospital and informed consent was obtained from the parent of each participant. CSF samples were received and remained deidentified for these studies.

### Cell Culture and Conditioned Media Preparation

LUHMES wild-type control, and MECP2 knock-out 2_7 cell line were differentiated and conditioned media was collected. Nunclon flasks and plates were treated with a 44 µg/ml Poly-L-Ornithine (Sigma P3655) and 1 µg/ml fibronectin (Sigma F1141) solution overnight in a 37°C incubator. LUHMES cells were differentiated as follows: three million cells were plated in a T75 flask with proliferation media (Advanced DMEM/F12 (Gibco 12634-010) with N2 (Gibco 17502048), 2mM L-glutamine (Sigma G7513), and 40ng/ml beta-FGF (R&D Systems 4114-TC-01M). After 24 hours, media was changed to differentiation media (Advanced DMEM/F12 with N2, 2mM L-glutamine, 1mM DbcAMP (Sigma D0627), 1 µg /ml tetracycline (Sigma T7660), and 2 ng/ml GDNF (R&D Systems 212-GD-050) for a pre-differentiation phase of two days. Pre-differentiated cells were lifted with trypsin method. Trypsin activity was blocked with aprotinin after lifting the cells. To reduce background signal in mass spectrometry, the last phase of differentiation utilized high purity and BSA-free components including high purity N2 components. A 100x high purity N2 solution was made with 10 mg/ml human holo-transferrin (Sigma T4132), 0.5mg/ml human recombinant insulin solution (Sigma I9278), 0.63 µg/ml progesterone (Sigma P6149), 1.61 mg/ml putrescine dihydrochloride (Sigma P5780), 0.52 µg/ml sodium selenite (Sigma S5261) and DMEM/F12 (Thermo Fisher 21331020). One million pre-differentiated cells were plated to each well of a Nunclon 6-well dish with 2 ml of the high purity differentiation media: DMEM/F12 (Thermo Fisher 21331020) containing high purity N2 (above), 2 mM L-glutamine (Sigma G7513), 1 mM DbcAMP (D0627), 1 µg/ml tetracycline (T7660), and 2 ng/ml GDNF (R&D Systems 212-GD-050). Cells conditioned the media for 3 days at 5% CO2 in a 37°C incubator. On the third day, the conditioned media was collected and Complete antiprotease (Roche 11697498001) was added. Cellular debris was pelleted at 16,000 x g in an Eppendorf microcentrifuge at 4°C for 20 minutes. The supernatant was collected and flash frozen on dry ice. A trichloroacetic acid (TCA) precipitation was done on 750 µl of the conditioned cell media by adding 9.8 µ g sodium deoxycholate per 100 µl of conditioned media followed by trichloroacetic acid to 10%. The solution was incubated on ice for 20 minutes to precipitate out proteins. The solution was centrifuged at 16,000 x g for 15min at 4°C. TCA supernatant was aspirated out and the pellet was washed in an equal volume of ice-cold acetone and vortexed. Precipitate was repelleted by centrifugation at 16,000 x g at 4°C for 10minutes. Acetone was aspirated and the pellet was lightly air-dried, dissolved in 200 µl of 8M Urea, and flash frozen on dry ice.

### Mass Spectrometry Emory

#### Sample Processing

All CSF (5μl) samples were diluted with 50 μl of 50 mM NH4HCO3 and treated with TCEP and CAA and heated at 90°C for 10 minutes. The samples were digested with 1:20 (w/w) lysyl endopeptidase (Wako) at 25°C overnight. Further overnight digestion was carried out with 1:20 (w/w) trypsin (Promega) at 25°C. Resulting peptides were desalted with a HLB microelution plate (Waters) and dried under vacuum.

#### Tandem Mass Tag (TMT) Labeling

For each sample, labeling was performed as previously described ^22,67^. Briefly, each was re-suspended in 100 mM TEAB buffer (100 μL). The TMT and TMTPro labeling reagents were equilibrated to room temperature, and anhydrous ACN (256 μL) was added to each reagent channel. Each channel was gently vortexed for 5 min, and then 41 μL from each TMT channel was transferred to the peptide solutions and allowed to incubate for 1 h at room temperature. The reaction was quenched with 5% (vol/vol) hydroxylamine (8 μl) (Pierce). All channels were then combined and dried by SpeedVac (LabConco) to approximately 150 μL and diluted with 1 mL of 0.1% (vol/vol) TFA, then acidified to a final concentration of 1% (vol/vol) FA and 0.1% (vol/vol) TFA. Peptides were desalted with a 30 mg C18 Sep-Pak column (Waters). Each Sep-Pak column was activated with 1 mL of methanol, washed with 1 mL of 50% (vol/vol) ACN, and equilibrated with 2×1 mL of 0.1% TFA. The samples were then loaded and each column was washed with 2×1 mL 0.1% (vol/vol) TFA, followed by 1 mL of 1% (vol/vol) FA. Elution was performed with 2 volumes of 0.5 mL 50% (vol/vol) ACN. The eluates were then dried to completeness.

#### High pH Fractionation

High pH fractionation was performed essentially as described ^68,69^ with slight modification. Dried samples were re-suspended in high pH loading buffer (0.07% vol/vol NH4OH, 0.045% vol/vol FA, 2% vol/vol ACN) and loaded onto Water’s BEH C18 column (2.1mm x 150 mm with 1.7 µm beads). An Thermo Vanquish system was used to carry out the fractionation. Solvent A consisted of 0.0175% (vol/vol) NH4OH, 0.01125% (vol/vol) FA, and 2% (vol/vol) ACN; solvent B consisted of 0.0175% (vol/vol) NH4OH, 0.01125% (vol/vol) FA, and 90% (vol/vol) ACN. The sample elution was performed over a 22 min gradient with a flow rate of 0.6 mL/min from 0 to 50% solvent B. A total of 96 individual equal volume fractions were collected across the gradient and subsequently pooled by concatenation into 48 fractions for the Mecp2 TMT batches. For the Cdlk5 batch, 192 fractions were collected and combined into 96 fractions. All fractions were dried to completeness using a vacuum centrifugation.

#### Liquid Chromatography Tandem Mass Spectrometry for TMT

Each of the peptide fractions was resuspended in loading buffer (0.1% FA, 0.03% TFA, 1% ACN). Peptide eluents were either separated on a self-packed C18 (1.9 µm Dr. Maisch, Germany) fused silica column (15 cm × 100μM internal diameter (ID), New Objective, Woburn, MA) or a Water’s 1.7 µm CSH C18 column (15 cm × 150μM internal diameter). An Easy nLC 1200 (ThermoFisher Scientific) or Ultimate U300 RSLCnano (Thermo Scientific) was used to elute the peptide ion. Mass spectra were collected either on a Fusion Lumos or Fusion Eclipse mass spectrometer. Both mass spectrometers were outfitted with the FAIMS Pro ion mobility source.

#### Liquid Chromatography Tandem Mass Spectrometry for Parallel Reaction Monitoring (PRM)

AQUA standard peptides (ThermoFisher Scientific) were spiked into digested mouse CSF samples. For each sample and equivalent of 1 ul of CSF was loaded onto a Water’s 1.7 µm CSH C18 column (15 cm × 150μM internal diameter). Peptides were eluted using a Ultimate 3000 RSLCnano and PRM spectra were collected using an Orbitrap HFX mass spectrometer.

#### Data Processing Protocol

All TMT raw files were searched using Thermo’s Proteome Discoverer suite (version 2.4.1) with Sequest HT. The spectra were searched against rat or mouse uniprot database. Search parameters included 20ppm precursor mass window, 0.05 Da product mass window, dynamic modifications methione (+15.995 Da), deamidated asparagine and glutamine (+0.984 Da), phosphorylated serine, threonine and tyrosine (+79.966 Da), and static modifications for carbamidomethyl cysteines (+57.021 Da) and N-terminal and Lysine-tagged TMT (+229.26340 Da or +304.207 Da). Percolator was used filter PSMs to 0.1%. Peptides were grouped using strict parsimony and only razor and unique peptides were used for protein level quantitation. Reporter ions were quantified from MS2 scans using an integration tolerance of 20 ppm with the most confident centroid setting. Only unique and razor (i.e., parsimonious) peptides were considered for quantification. PRM spectra were processed using the Skyline quantitation suite ^70^.

### Mass Spectrometry Ann Arbor

#### Sample Preparation

For tissues, samples were washed twice in 1X PBS and lysed in 8M urea, 50mM Tris HCl, pH 8.0, 1X Roche Complete Protease Inhibitor and 1X Roche PhosStop. Other samples were processed directly for protein quantification using Qubit fluorometry following by digestion overnight with trypsin. Briefly, samples were reduced for 1h at RT in 12mM DTT followed by alkylation for 1h at RT in 15mM iodoacetamide. Trypsin was added to an enzyme:substrate ratio of 1:20. Each sample was acidified in formic acid and subjected to SPE on an Empore SD C18 plate. For TMT labeling, after trypsin digestion ach sample was acidified in formic acid and subjected to SPE on an Empore SD C18 plate (3M catalogue# 6015 SD). Each sample was lyophilized and reconstituted in 140mM HEPES, pH 8.0, 30% acetonitrile.

#### Label Free Quantification Mass Spectrometry

A 2μg aliquot was analyzed by nano LC/MS/MS with a Waters NanoAcquity HPLC system interfaced to a ThermoFisher Fusion Lumos. Peptides were loaded on a trapping column and eluted over a 75μm analytical column at 350nL/min; both columns were packed with Luna C18 resin (Phenomenex). A 4h gradient was employed. The mass spectrometer was operated in data-dependent mode, with MS and MS/MS performed in the Orbitrap at 60,000 FWHM resolution and 15,000 FWHM resolution, respectively. APD was turned on. The instrument was run with a 3s cycle for MS and MS/MS. The acquisition order was randomized. Data Processing Data were processed through the MaxQuant software v1.6.2.3 (www.maxquant.org). Data were searched using Andromeda with the following parameters: Enzyme: Trypsin, Database: Uniprot Rat, Fixed modification: Carbamidomethyl (C), Variable modifications: Oxidation (M), Acetyl (Protein N-term), Fragment Mass Tolerance: 20ppm Pertinent. MaxQuant settings were: Peptide FDR 0.01 Protein FDR 0.01 Min. peptide Length 7 Min. razor + unique peptides 1 Min. unique peptides 0 Min. ratio count for LFQ 1 Second Peptides^ TRUE Match Between Runs* TRUE

#### TMT Quantification Mass Spectrometry

40μL of acetonitrile was added to each TMT tag tube and mixed aggressively. Tags were incubated at RT for 15min. 30μL of label was added to each peptide sample and mixed aggressively. Samples were incubated in an Eppendorf Thermomixer at 300rpm 25°C for 1.5h. Reactions were terminated with the addition of 8μL of fresh 5% hydroxylamine solution and 15min incubation at room temperature. Samples were subjected to high pH reverse phase fractionation as follows; Buffers: Buffer A: 10mM NaOH, pH 10.5, in water Buffer B: 10mM NaOH, pH 10.5, in acetonitrile. We used XBridge C18 colums, 2.1mm ID x 150mm length, 3.5μm particle size (Waters, part #186003023) attached to a Agilent 1100 HPLC system equipped with a 150μL sample loop operating at 0.3mL/min, detector set at 214 nm wavelength. Dried peptides were resolubilized in 150μL of Buffer A and injected manually. Fractions were collected every 30s from 1-49min (96 fractions total, 150μL/fraction). We analyzed by mass spectrometry 10% per pool for the full proteome in a nano LC/MS/MS with a Waters NanoAcquity HPLC system interfaced to a ThermoFisher Fusion Lumos mass spectrometer. Peptides were loaded on a trapping column and eluted over a 75μm analytical column at 350nL/min; both columns were packed with Luna C18 resin (Phenomenex). Each high pH RP fraction was separated over a 2h gradient (24h instrument time total). The mass spectrometer was operated in data-dependent mode, with MS and MS/MS performed in the Orbitrap at 60,000 FWHM resolution and 50,000 FWHM resolution, respectively. A 3s cycle time was employed for all steps. Data Processing Data were processed through the MaxQuant software v1.6.2.3 (www.maxquant.org). Data were searched using Andromeda with the following parameters: Enzyme: Trypsin Database: Uniprot Rat, Fixed modification: Carbamidomethyl (C) Variable modifications: Oxidation (M), Acetyl (Protein N-term), Phopho (STY; PO4 data only). Fragment Mass Tolerance: 20ppm. Pertinent MaxQuant settings were: Peptide FDR 0.01 Protein FDR 0.01 Min. peptide Length 7 Min. razor + unique peptides 1 Min. unique peptides 0 Second Peptides FALSE Match Between Runs FALSE The protein Groups.txt files were uploaded to Perseus v1.5.5.3 for data processing and analysis.

#### AQUA Mass Spectrometry

Synthetic peptides labeled with Arginine (13C6,15N4) or Lysine (13C6,15N2) at >95% purity were made by New England Peptide MA 01440 USA. The following peptides were used: Myh9 AGVLAHLEEER; IAQLEEQLDNETK. Pon1 IFFYDSENPPGSEVLR; LLIGTVFHR. App TEEISEVK; THTHIVIPYR. A2m AIAYLNTGYQR; LPSDVVEESAR. Apoa1 DYVSQFESSTLGK; WNEEVEAYR. Apob TEVIPPLIENR; GFEPTLEALFGK. C3 GLEVSITAR; SSVAVPYVIVPLK. C9 SIEVFGQFQGK; TTSFNANLALK. Thbs1 FVFGTTPEDILR; IENANLIPPVPDDK. A 3-4 μg aliquot of each CSF tryptic peptide digests was spiked with isotopologe peptides at a concentration of 100 or 133 fmol/μg peptide digest. Peptides mixes were analyzed in analytical duplicate by nano LC/PRM using a Waters NanoAcquity HPLC system interfaced to a ThermoFisher Fusion Lumos mass spectrometer. 1.5μg per sample was loaded on a trapping column and eluted over a 75μm analytical column at 350nL/min; both columns were packed with Luna C18 resin (Phenomenex). A 1h gradient was employed. The mass spectrometer was operated in PRM mode without scheduling; instrument settings included 15,000 FWHM resolution, NCE 30, AGC target value 5e4, and maximum IT of 22ms. Data were processed using Skyline v4.2.

### Data Processing

Proteomics data were log2 converted. Data analysis was performed with two methods. We used Qlucore Omics Explorer Version 3.6(33) normalizing data to a mean of 0 and a variance of 1. No filtering by standard deviation was applied. All data were thresholded by a log2 fold of change of 0.5 and a non-corrected p value of 0.05.

A second method used was OmicLearn (v1.0.0) for performing the data analysis, model execution, and generating the plots and charts ^42^. Machine learning was done in Python (3.8.8). Feature tables were imported via the Pandas package (1.0.1) and manipulated using the Numpy package (1.18.1). The machine learning pipeline was employed using the scikit-learn package (0.22.1). For generating the plots and charts, Plotly (4.9.0) library was used. No normalization on the data was performed. To impute missing values, a Mean-imputation strategy is used. Features were selected using a ExtraTrees (n_trees=100) strategy with the maximum number of 20 features. Normalization and feature selection was individually performed using the training data of each split. For classification, we used either a XGBoost-Classifier (random_state = 23 learning_rate = 0.3 min_split_loss = 0 max_depth = 6 min_child_weight = 1), a AdaBoost-Classifier (random_state = 23 n_estimators = 100 learning_rate = 1.0), or RandomForest-Classifier (random_state = 23 n_estimators = 100 criterion = gini max_features = auto). Clasifiers were chosen based on the proximity of ROC curves to a value of 1. When using (RepeatedStratifiedKFold) a repeated (n_repeats=10), stratified cross-validation (n_splits=5) approach to classify datasets based on their genotype.

### RNAseq and Single Cell RNAseq

RNAseq data analysis was described in Wynne et al. ^71^. Single cell RNA seq data were described in ^72^. Gene expression data matrix (matrix.csv) and cell metadata (metadata.csv) data were downloaded from <u>the Allen Institute Portal</u> and processed as described ^71^ with the Qlucore Omics Explorer Version 3.6(33). Data were log2 converted and normalized to a mean of 0 and a variance of 1. 2D t-SNE plots were generated using a perplexity of 40 and default settings.

### Bioinformatic Analyses

Gene ontology analyses were performed with Cluego and HumanBase ^47^. ClueGo v2.58 run on Cytoscape v3.8.2 ^46,73^. ClueGo was run querying GO CC, REACTOME, KEGG and WikiPathways considering all evidence at a Medium Level of Network Specificity and selecting pathways with a Bonferroni corrected p value <10E-3. ClueGo was run with Go Term Fusion. HumanBase was run using default webased parameters^47^. *In silico* interactome data were downloaded from Genemania predicted and physical interactions and processed in Cytoscape v3.8.2 ^73^. Interactome connectivity graph parameters were generated in Cytoscape.

### Statistical Analyses

Volcano plot p values were calculated using Qlucore Omics Explorer Version 3.6(33) without multiple corrections. Experiments in Figure 4A-B and G statistical analyses were performed with the engine https://www.estimationstats.com/#/ with a two-sided permutation t-test and alpha of 0.05 ^74^. ROC analysis and paired t-test were performed with Prism v9.2.0(283).

### Data Availability

The mass spectrometry proteomics data have been deposited to the ProteomeXchange Consortium via the PRIDE ^75^ partner repository with dataset identifiers: PXD029808, PXD029809, PXD029811, PXD029835

## Acknowledgements

VF was funded by the Rett Syndrome Research Trust, the Loulou Foundation, and 1RF1AG060285. S Rangaraju was partly funded by the NIH (5R01NS114130). S Rayaprolu was partly supported by the NIH (F32 AG064862). SC was funded by the Rett Syndrome Research Trust and Simons Initiative for the Developing Brain

## Figure Legends

**Fig. S1.**
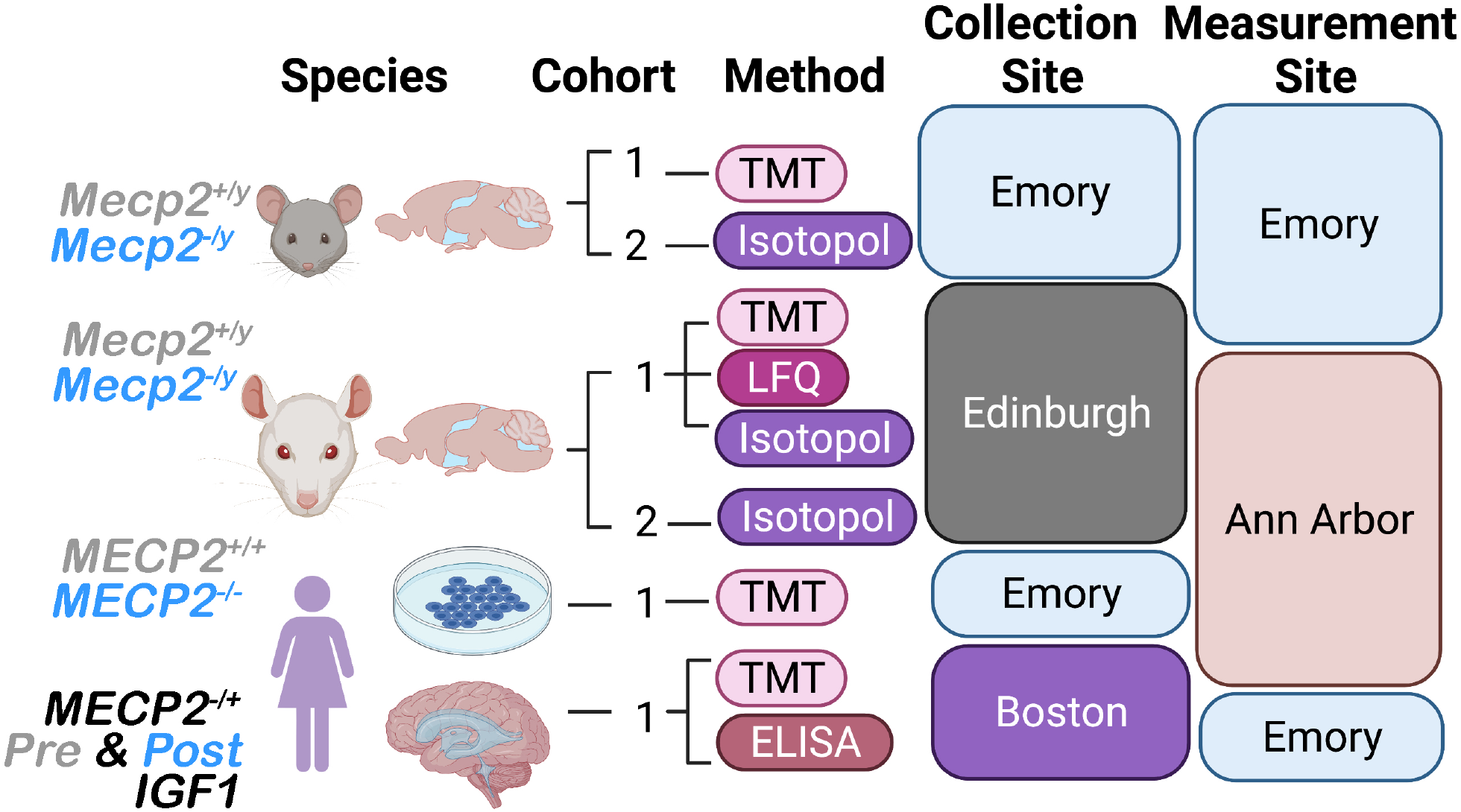
Design of a Robust Strategy to Identify Consensus Rett Syndrome Proteomes for Ontology Inference and Biomarker Selection. Diagram shows three species and experimental systems. Cohorts represent independent collections of samples with the strategy used for analyte quantification, the place of sample collection, and sample measurement location. Isotopol refers to AQUA and modified AQUA strategies, LFQ corresponds to label free quantification, and TMT denotes Tandem Mass Tagging.

**Fig. S2.**
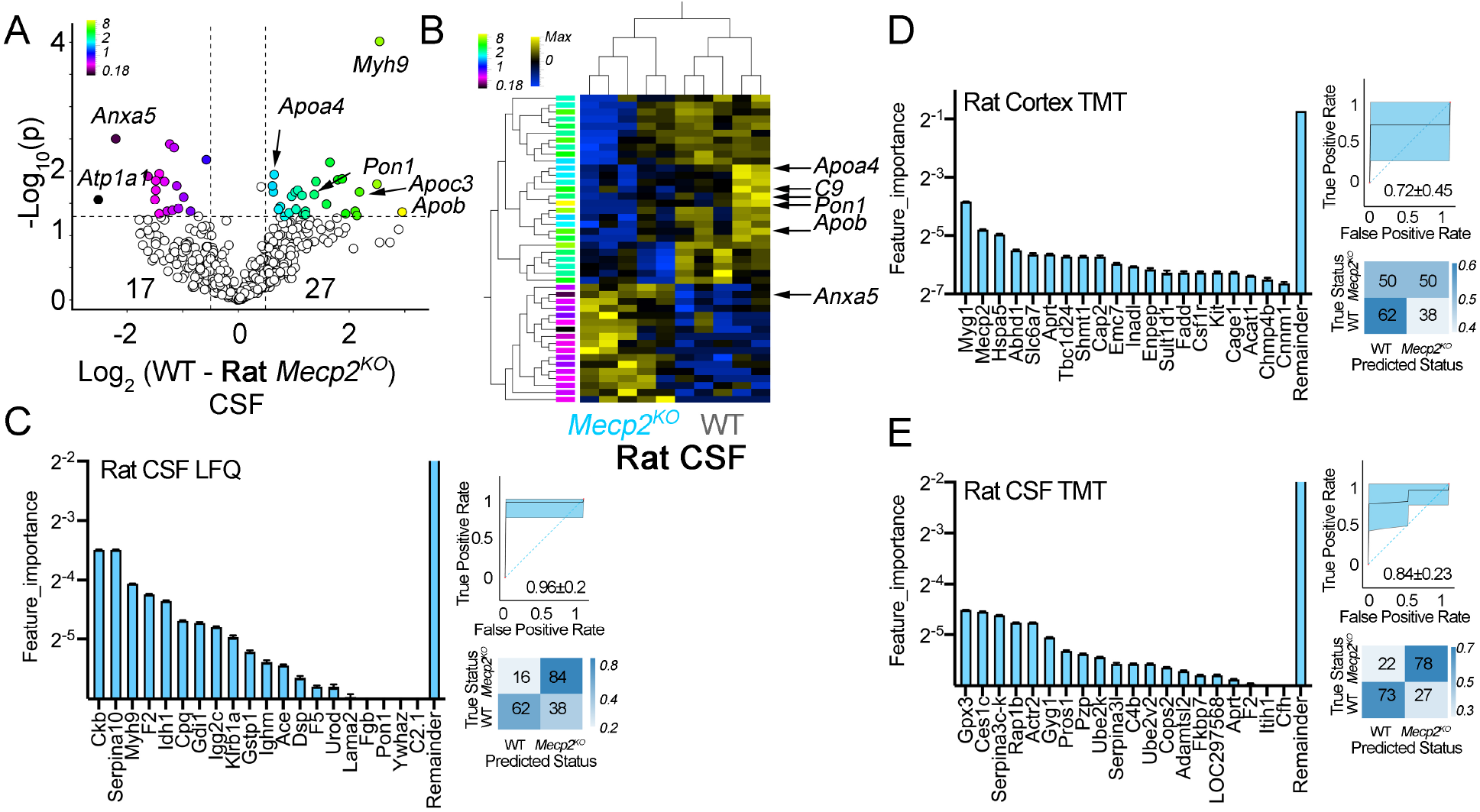
Orthogonal Approaches to Identify Rett Syndrome CSF Proteomes. **A** volcano plot of LFQ mass spectrometry determinations in rat cerebrospinal fluid. Cutoffs at log2 0.5 fold of change in protein abundance and a p value <0.05, n=5. See table SS2. **B** shows clustered heat maps of hits selected in A. For A and B, see legend to Figure 1 for additional details. **C** to **E**, depict analysis of the indicated mass spectrometry datasets using a Random Forest machine learning algorithm. Bar graph showing main hits discriminating wild type and *Mecp2* mutant CSF in the decision tree. Inserts show performance of the machine learning protocol estimated by ROC analysis and confusion matrix.

**Fig. S3.**
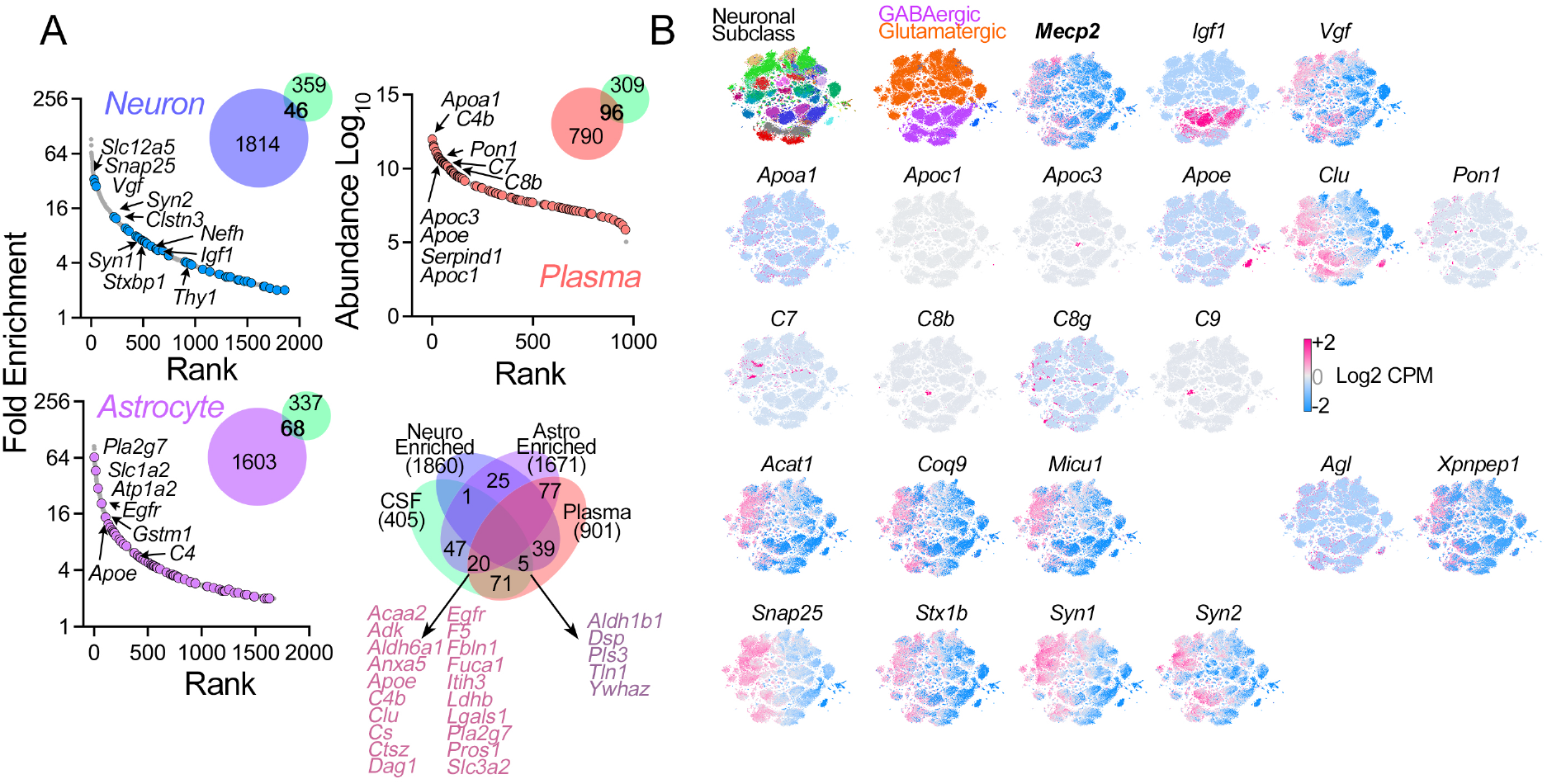
Expression Patterns of Ontology Selected Analytes in Brain Cells and Plasma. **A.** Fold enrichment and rank order of mRNAs most expressed in neurons and astrocytes according to Zhang ^76,77^ or abundance in the plasma proteome according to Geyer et al ^78^. Superimposed are CSF hits. Venn diagrams present overlaps with each cell type gene expression category or plasma proteome. **B** depicts a t-SNE cell atlas generated with the expression levels of all transcripts encoding selected hits from the mouse CSF Mecp2-sensitive proteome. The t-SNE atlas encompasses >20 areas of mouse cortex and hippocampus, totaling 76,307 cells ^79^. Color codes denote neuronal subclasses described by Yao et al. ^79^. Neurotransmitter annotation is depicted as well as the expression levels of Mecp2 mRNA across brain regions and cell types. Each atlas depicts the mRNA expression of the indicated analyte. Note analytes such as Apoc1 whose mRNA is not detectable in this dataset. t-SNE cell atlases were assembled using the Allen single-cell RNAseq dataset as describe by Wynne et al ^80^.

**Fig. S4.**
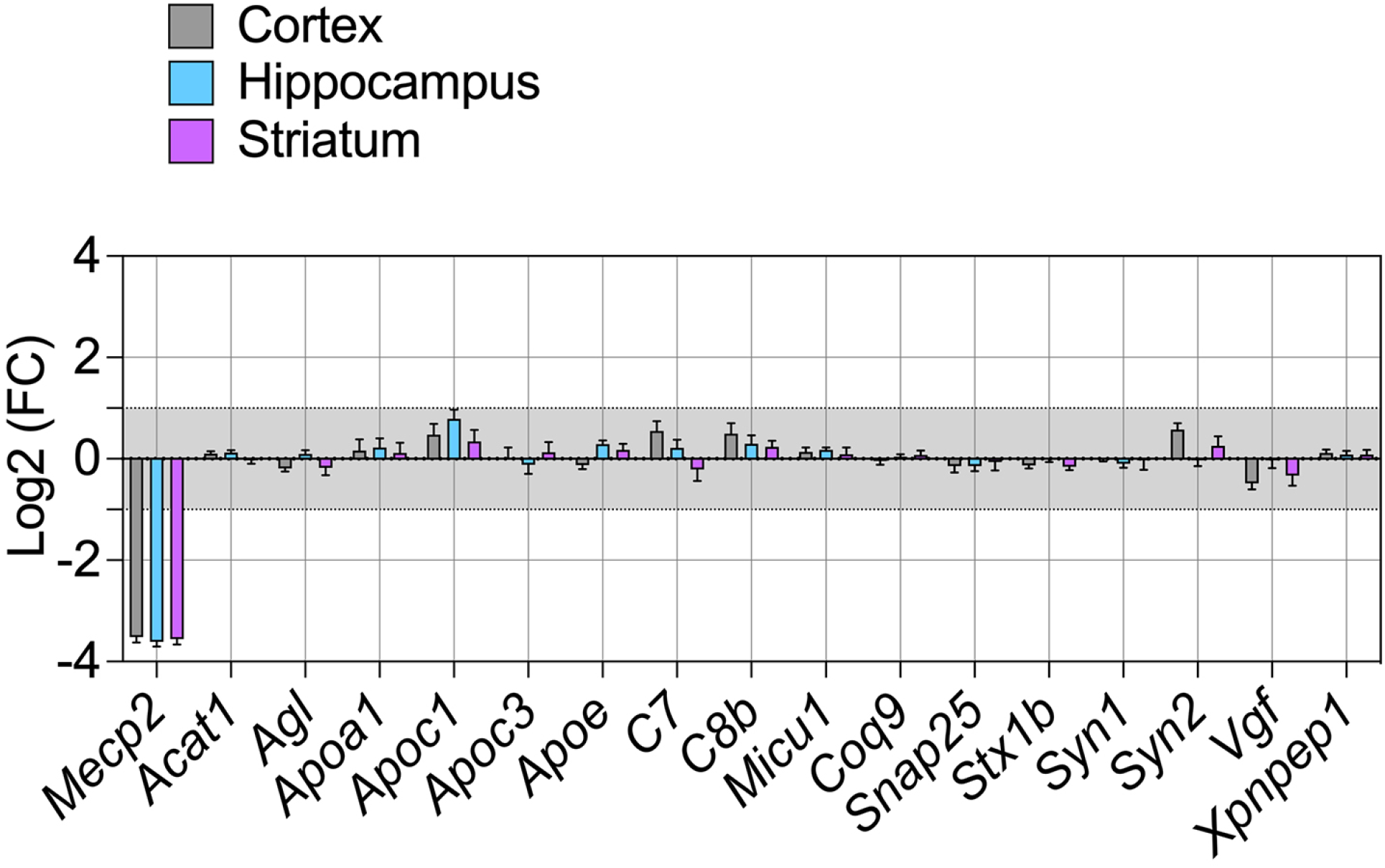
Brain Parenchyma Transcripts Levels of Analytes Selected by Ontology and with Biomarker Potential. RNAseq analysis of transcript expression of indicated analytes in three microdissected brain regions from wild type and Mecp2 mutant brains. Mecp2 mRNA are presented as a control. N=5 animals per each phenotype. Only Mecp2 mRNA is differentially expressed p for three brain regions p< 1.40E-236, Benjamini and Hochberg corrected Wald test. See table SS4.

